# Faces capture spatial attention only when we want them to: an inattentional blindness EEG study

**DOI:** 10.1101/2023.05.17.541247

**Authors:** Zeguo Qiu, Xue Lei, Stefanie I. Becker, Alan J. Pegna

**Author notes:** **Correspondence** Zeguo Qiu, School of Psychology, The University of Queensland, Brisbane 4072, Australia. **E-mail addresses:** Zeguo Qiu; Xue Lei; Stefanie I. Becker; Alan J. Pegna.

## Abstract

Previous research on emotional face processing has shown that emotional faces such as fearful faces may be processed without visual awareness. However, evidence for nonconscious attention capture by fearful faces is limited. In fact, studies using sensory manipulation of awareness (e.g., backward masking paradigms) have shown that fearful faces do not attract attention during subliminal viewings nor when they were task-irrelevant. Here, we used a three-phase inattentional blindness paradigm and electroencephalography to examine whether faces (fearful and neutral) capture attention under different conditions of awareness and task-relevancy. We found that the electrophysiological marker for attention capture, the N2-posterior-contralateral (N2pc), was elicited by face stimuli only when participants were aware of the faces and when they were task-relevant (phase 3). When participants were unaware of the presence of faces (phase 1) or when the faces were irrelevant to the task (phase 2), no N2pc was observed. Together with our previous work, we concluded that fearful faces, or faces in general, do not attract attention unless we want them to.

## Introduction

Human faces have been found to be prioritised over non-face information (Reynolds & Roth, 2018; Zhou et al., 2021), possibly because faces convey important social information including a person’s emotional states. The emotional expression of faces facilitates our social interactions with others and provides information about our environment. For example, being able to recognise fearful expressions may be crucial for us to quickly respond to immediate danger in our surrounds.

Using a variety of experimental paradigms, it has been shown that emotional faces can be detected faster than their neutral counterparts (Frischen et al., 2008). When emotional faces are presented as distractors or task-irrelevant stimuli, they can also capture participants’ attention, slowing down the response to the target (Elam et al., 2010; Fox et al., 2002; Glickman & Lamy, 2018) or lowering task accuracy (Grose-Fifer et al., 2013). Consistent with these behavioural observations, neuroimaging studies have shown that a target that is validly cued by an emotional face is associated with an enhanced neural response, compared to a target that does not follow an emotional face (Holmes et al., 2009; Pourtois & Vuilleumier, 2006; Qiu et al., 2023a). In a similar vein, studies on patients with affective blindsight (for a review see Celeghin et al., 2015) and patients with hemianopia (for a review see Làdavas & Bertini, 2021) have suggested that emotional faces may be processed in the absence of visual awareness.

In healthy populations, some research has similarly shown that there may be subliminal or nonconscious processing of emotional faces (Del Zotto & Pegna, 2015; Kiss & Eimer, 2008; Pegna et al., 2008; Pegna et al., 2011; Suzuki & Noguchi, 2013). However, evidence for nonconscious *attention capture* by emotional faces (i.e., their ability to attract attention) is extremely limited. In fact, using masking techniques, a series of studies have shown that attention capture by emotional faces is possible only when participants are aware of the stimuli (Hedger et al., 2019; Qiu et al., 2022a, 2023b; Tipura & Pegna, 2022).

One effective way to measure attention capture is by examining its electrophysiological marker with electroencephalography (EEG). Specifically, our previous work has shown that a neural marker for attention capture, the N2-posterior-contralateral (N2pc), to a fearful face in a face pair can be found only during supraliminal viewings of the faces (Qiu et al., 2022a, 2023a, 2023b). When the faces were presented very briefly (16ms) and immediately backward masked, no N2pc was observed in the data. These findings were taken to suggest that attentional capture by fearful faces requires visual awareness. Furthermore, and importantly, when participants were required to attend to non-facial features superimposed on the face images, the task-irrelevant fearful faces did not capture attention even when they were clearly visible to the participants (Qiu et al., 2022a, 2023b). Thus, it seems that only supraliminal, task-relevant fearful faces attract attention and elicit a significant N2pc.

However, masking techniques are by no means the only way of investigating nonconscious processing of visual stimuli. As pointed out by Diano and colleagues (2017), two main types of experimental paradigms exist in awareness research: sensory paradigms such as backward masking, and attentional paradigms including the attentional blink and inattentional blindness (IB) paradigms. It was proposed that different neural mechanisms could subserve the nonconscious processing of emotional stimuli in different paradigms (Diano et al., 2017). Indeed, in our recent meta-analysis of functional magnetic resonance imaging studies on nonconscious emotion processing, we found that different paradigms were associated with activations of largely distinct brain regions during the processing of unseen (for sensory paradigms) and unattended (for attentional paradigms) emotional faces (Qiu et al., 2022c). Hence, one important question is whether we can replicate our previous findings regarding the N2pc to emotional faces with an attentional paradigm, or whether our findings are restricted to sensory paradigms.

In a recent study, we used a rapid serial visual presentation paradigm to investigate whether and how the N2pc to a fearful face could be modulated by different levels of awareness of the targets induced by the attentional blink (Qiu et al., 2022b). We found a significant N2pc to the fearful target face even in conditions where visual awareness was most strongly impeded, although the N2pc was substantially diminished compared to other conditions. However, in attentional paradigms such as the attentional blink, awareness may not be fully impeded and some residual attention may be present for conscious processing of the facial expressions.

An alternative method to investigate inattention and awareness is the three-phase IB paradigm. In this paradigm, stimuli are usually kept constant throughout the experiment, and awareness is manipulated through task instructions. Specifically, in the first phase, participants are not informed about the presence of a critical stimulus (e.g., a face), and they are asked to perform a task on non-critical stimuli (e.g., non-face shapes). In the second phase, participants are told that there is in fact a critical stimulus in some trials, but they are asked to perform the task on non-critical stimuli only and not to respond to the critical stimulus. In the third and final phase, participants are asked to additionally respond to the critical stimulus. Thus, in a typical IB study, it is possible to disentangle effects of awareness and task-relevancy of the stimuli: the intended difference between phase 1 and 2 is whether the participants are aware of the critical stimulus (phase 2) or not (phase 1), whereas the intended difference between phase 2 and 3 is whether the now-aware critical stimulus is task-relevant (phase 3) or not (phase 2).

The IB paradigm has been widely used in the awareness literature (e.g., Harris et al., 2020; Pitts et al., 2018; Tsuchiya et al., 2015). However, to the best of our knowledge, attention capture by fearful faces has not been examined with this IB paradigm. The aim of the present study was to use this paradigm to address some of the limitations in our previous work. Specifically, in real life, stimuli are rarely flashed for an extremely short duration (i.e., 16ms) and masked. Rather, we are constantly presented with streams of visual inputs. Attention may in fact be the key to determine if a stimulus has access to conscious awareness in this case. That is, it remains unclear whether clearly visible faces, emotional faces in particular, can capture spatial attention (by eliciting an N2pc) when we are unaware of them during inattentional blindness. Additionally, as mentioned, in our previous attentional blink paradigm, it was perhaps impossible to impede visual awareness sufficiently as stimuli were always presented with some temporal separation (Qiu et al., 2022b). The time lag between the two target pairs in the attentional blink experiment could be sufficient for attention to be captured by the fearful face. Here, we aimed to implement complete inattention to face images in phase 1 of the IB procedure.

Furthermore, one important advantage of the IB paradigm is that the stimuli are kept constant throughout the experiment and we can directly compare effects of awareness and effects of task-relevancy. This can be done by examining modulations related to awareness (i.e., phase 1 vs. 2) and those related to task-relevancy (i.e., phase 2 vs. 3) on the awareness-related components; specifically by examining the visual awareness negativity (VAN) and the late positivity (LP). The VAN is a negativity for aware stimuli in the event-related potential (ERP) signals which appears at around 200ms post stimulus onset, and it is most pronounced at posterior brain regions. The LP is a sustained positivity towards aware stimuli that appears after 300ms post stimulus at parietal regions. These two awareness-related components have been found to correlate with awareness (for reviews see Cohen et al., 2020; Förster et al., 2020), and the VAN has further been suggested to be the first and more reliable neural marker for perceptual awareness, including but not limited to visual awareness (Dembski et al., 2021). Moreover, our previous work has shown that task-relevancy of the stimuli can further enhance these components (Qiu et al., 2023a), suggesting that the VAN and the LP can be modulated by participants’ attention.

To summarise, in this study, we focused on three ERPs: the N2pc that indicates spatial attention capture, and the VAN and the LP that index visual awareness. We firstly hypothesised that the face stimuli would only attract attention when they were consciously perceived and task-relevant, consistent with our previous findings (Qiu et al., 2022a). Operationally, we expected to find an N2pc to the faces only in phase 3 of the IB experiment. Secondly, we predicted that a fearful face would elicit a larger N2pc, if observed, than a neutral face, in alignment with previous literature on emotional attention capture. Lastly, we expected to find significant VAN and LP when comparing phase 2 and phase 1, reflecting the effects of awareness. We also expected to find significant VAN and LP when comparing phase 3 and phase 2, which should reflect larger VAN and LP when the stimuli were task-relevant, in line with our previous findings (Qiu et al., 2023a).

## Method

### Participants

We determined the sample size with MorePower (Campbell & Thompson, 2012) using an effect size reported in a previous study with a similar design (Cohen ’s *d* = 0.24; Harris et al., 2020). A minimum of 22 participants were required for significant differences between the three phases in our 3(phase: phase 1, phase 2, phase 3) x 2(face emotion: fearful, neutral) x 2(laterality: contralateral, ipsilateral) design, with 90% power and a two-tailed alpha level of .05. We recruited 38 participants and compensated them with either monetary reimbursement or course credits. Among the 38 people, 13 participants reported seeing the face images after phase 1 and were labelled as “noticers”, and their data were not analysed in the current study. The remaining 25 participants were included in the main “attentionally blind” group. Data from one “attentionally blind” participant were excluded because of a 0 accuracy at the face task in phase 3. Therefore, the final sample constituted 24 participants (*M*_*age*_ = 22.7 years, *SD*_*age*_ = 3.1 years, 4 males, 20 females).

This study was approved by the University of Queensland ethics committee.

### Apparatus and experimental stimuli

All stimuli were presented on a 24-inch ASUS LCD monitor (resolution: 1920 × 1080 pixels) which was placed 70 cm away from the participant’s eyes. The program was run in PsychoPy3 (Peirce et al., 2019).

Face images were obtained and adapted from the Radboud Faces Database (Langner et al., 2010). Images from a total of 16 models (8 males and 8 females) were used. Each face image was converted to greyscale and cropped into an oval or a rectangle shape (2.8° × 3.6° in visual angle; see Figure 1a). We used a scrambled filter tool (http://telegraphics.com.au/sw/product/scramble) to make target rectangles and distractor ovals (all 2.8° × 3.6° in visual angle). Specifically, each cropped face image was sliced into small squares (208 for oval shapes, 252 for rectangle shapes) and randomly re-assembled (see Figure 1a). Only the rectangles were task-relevant while the ovals were task-irrelevant. All photo editing was done in Photoshop 2021 version 22.4.0 (Adobe Systems, San Jose, CA).

**Figure 1.**
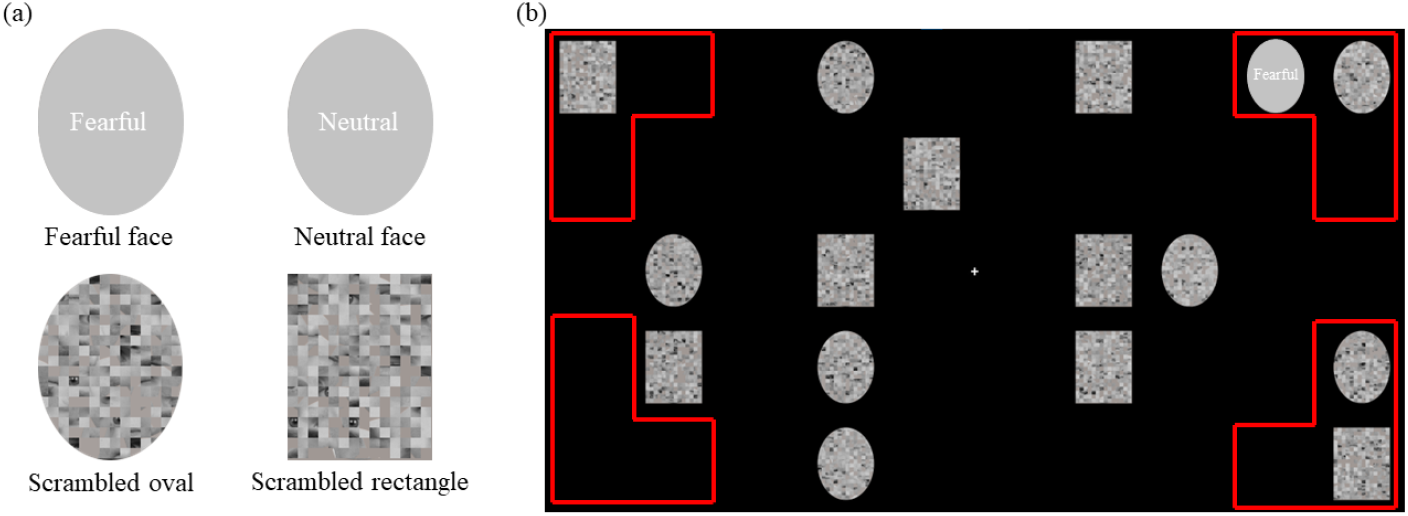
(a) Examples of experimental stimuli. (b) An example of the stimulus array in a given trial. The red lines highlight the possible locations of the single face image in a critical face-present trial. Note: The face stimuli are covered and de-identified in this picture in accordance with bioRxiv policies. Actual face images were used in the experiment.

Each target display consisted of 16 images randomly presented at 16 possible image positions in a 5-by-10 grid, with four rectangles and four ovals on each side of the screen. In the Critical trials (200 trials in each phase), one of the ovals was replaced by a face image and its location was restricted to one of 12 possible image positions at the four corners of the screen (three at each corner; see Figure 1b). The face image appeared equiprobably on either the attended or the unattended side, and it could either be a fearful face or a neutral face. We restricted the location of the face images in order to prevent the face images from being presented in participants’ foveal visual field and hence being noticed easily. In the Non-critical trials (50 trials in each phase), no face image was presented.

### Procedure

As shown in Figure 2, participants were first presented with a blank screen for a randomly selected duration between 500-800ms. Then, an arrow cue (either cueing left or right) was presented at the screen centre for 1000ms, followed by the target display, where a total of 16 images were presented on the screen for 400ms. Participants were required to memorise the locations of the four target rectangles on the cued screen side, while keeping their gaze fixated at a cross at the screen centre. Afterwards, participants saw a blank screen of 400ms and then a response map (a map of 5-by-10 letter “X”s). They were required to use the mouse to click on the Xs to indicate the spatial locations of the target rectangles in the previous screen (the rectangle localisation task). If a selection was correct, the X would turn green otherwise it would turn red. After selection of four Xs, participants needed to click the “submit” text at the screen centre to proceed to the next trial. Alternatively, if a fifth click on the screen was made, the trial would automatically end and the next trial would begin.

**Figure 2.**
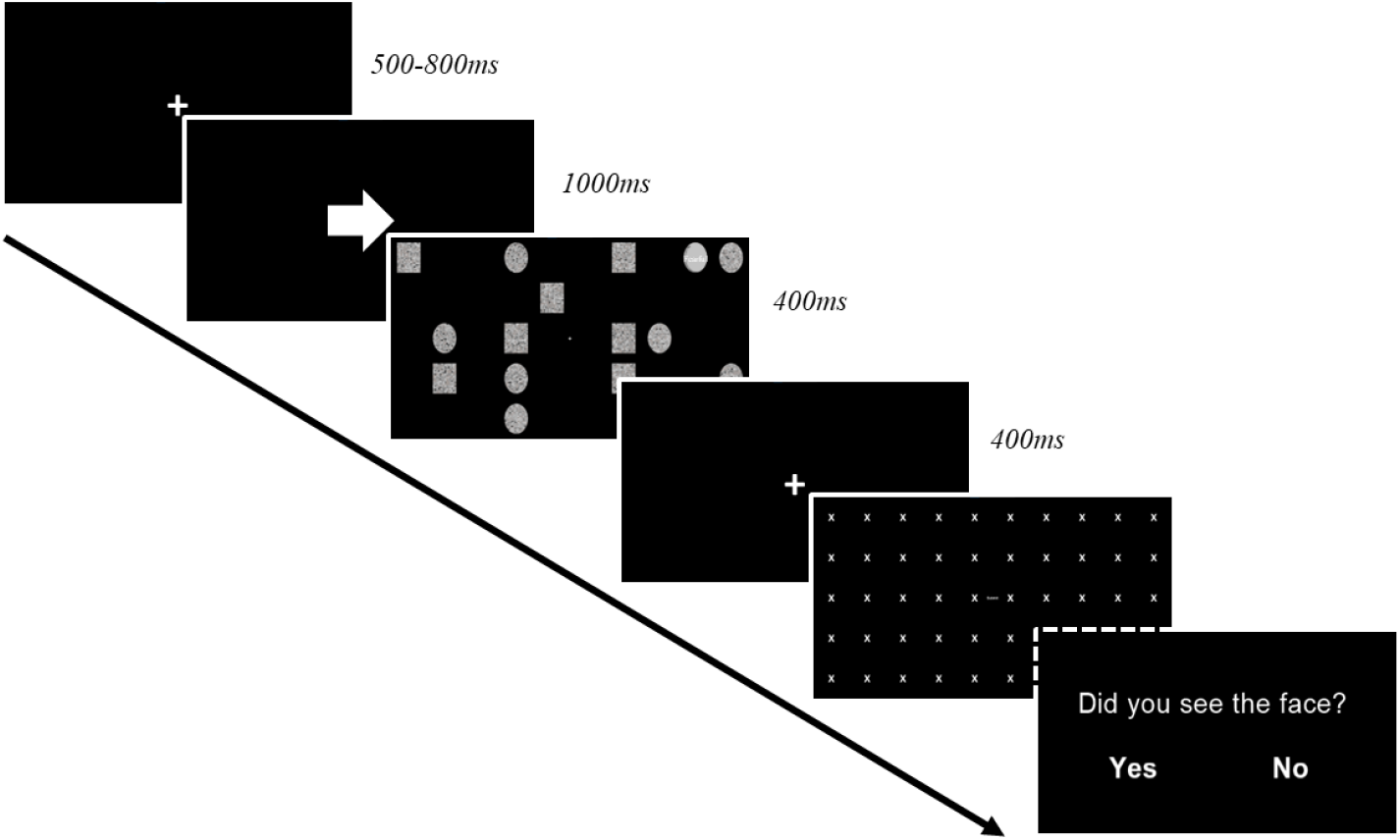
The full procedure of a trial. The question “Did you see the face?” was only presented in phase 3 of the experiment.

There were three phases in the experiment with five blocks of 50 trials in each phase. The order of critical and non-critical trials were randomised. In phase 1 (blocks one to five), participants were not informed about the presence of the face images. After phase 1, the experimenter asked the participants whether they had noticed anything strange or unexpected in the stimuli, and whether they had seen anything that was neither an oval nor a rectangular scrambled image. If participants responded yes to either question and reported seeing faces or face-like images, they were labelled as “noticers’, otherwise, they were labelled as “attentionally blind” to the critical face images. In phase 2 (blocks six to ten), participants were informed about the possible presence of face images, but continued to perform the rectangle localisation task and did not need to respond to the face images. In phase 3 (blocks 11 to 15), participants were required to perform the rectangle localisation task and then indicate whether they had seen a face in that trial. We did not require the participants to detect the emotion of the faces. Response to the second question was provided by using the mouse to click on the “Yes” or “No” text on the screen (Figure 2).

Twelve non-critical practice trials were provided before the actual experiment. Breaks were allowed between blocks and participants were instructed to keep fixated at the screen centre unless they had to move their eyes to select the target locations with the mouse.

### EEG recording and data pre-processing

Continuous EEG was recorded at 1024 Hz using the BioSemi ActiveTwo 64-electrode system (Biosemi, Amsterdam, Netherlands). We recorded the horizontal electrooculogram (EOG) using two bipolar electrodes placed laterally on the outer canthi of the eyes. The vertical EOG was recorded with an external electrode placed below participants’ left eye, paired with FP1. Recordings were referenced online to the CMS/DRL electrodes.

Pre-processing of the EEG data was performed with EEGLAB (Delorme & Makeig, 2004) and ERPLAB (Lopez-Calderon & Luck, 2014). We interpolated electrodes that produced noise throughout the experiment. Signals were re-sampled to 512 Hz offline, filtered from 0.1 to 30 Hz and notch-filtered at 50 Hz to remove line noise. All signals were then re-referenced to the average of all electrodes. Signals were segmented into epochs with a time window of 800ms from the onset of the target display, and baseline corrected using the interval of -100 to 0ms. Independent component analysis was performed on the epoched data to identify and remove eye-blink and eye-movement artefacts in the EEG. After eye-artefact removal, epochs were visually inspected trial by trial, and we removed epochs that contained minor eye-movements (using the horizontal EOG) or other artefacts. On average, 88% epochs were kept across participants.

To remove any practice effect across the three phases, we subtracted the average ERPs of the non-critical trials from the average ERPs of the critical trials for each phase. Analyses were then performed on the subtracted signals.

### Data analysis

#### Behavioural task performance

For the spatial localisation task, we compared the task accuracy (percent correct) across three phases with a one-way repeated-measures ANOVA.

For the face task in phase 3, we first compared the accuracy (percent correct) between two face emotion with a paired-samples *t*-test. We also applied the signal detection theory to the face task responses (Donaldson, 1992). Specifically, we calculated the d-primes (d’) and criterions (c) separately for fearful face and neutral face conditions. The d’ is an open-ended measure ranging from 0 to 2 and above with a larger d’ indicating high discriminability for the targets. A c around 0 indicates no response bias whereas a c above 0 indicates a response bias towards reporting target-absent, in our case not seeing a face, and a c below 0 indicates a bias towards reporting target-present, or seeing a face.

#### Mass univariate analysis

ERP analyses were conducted using the factorial mass univariate toolbox (Fields & Kuperberg, 2020) and the mass univariate toolbox (Groppe et al., 2011). We used the cluster-based permutation tests for significance testing correcting for multiple comparisons (2000 permutations). For cluster formation, the threshold was set at an alpha level of .05, and channels were considered as spatial neighbours if they were within 3.7cm of each other (Mean number of spatial neighbours: 2.9). For statistical significance, we used a family-wise alpha level of .05. Follow-up tests for any significant omnibus effect were conducted using the cluster-based permutation *t*-tests (2000 permutations).

To examine whether the critical face stimulus captured attention and hence elicited an N2pc, we calculated the ERPs contralateral and ipsilateral to the face images, collapsing across hemispheres, and conducted a 3(phase: phase 1, phase 2, phase 3) x 2(laterality: contralateral, ipsilateral) x 2(face emotion: fearful, neutral) repeated-measures ANOVA over the data.

## Results

### Behavioural task performance

A one-way repeated-measures ANOVA computed over the spatial localisation task performance (percentage of correct responses) revealed a significant main effect of phase, *F*(2,46) = 8.15, *p* = .001, *η*_*p*_^*2*^ = 0.26. Follow-up tests using Bonferroni correction for multiple comparisons showed that task accuracy was significantly higher in phase 2 (*M* = 0.83, *SD* = 0.10), compared to phase 1 (*M* = 0.79, *SD* = 0.11), *p* = .001, and phase 3 (*M* = 0.80, *SD* = 0.11), *p* = .007. There was no significant difference between task accuracies in phase 1 and 3, *p* = 1. The increased accuracy in phase 2 compared to phase 1 may indicate a practice effect. The decrease in accuracy in phase 3 compared to phase 2 is likely due to a higher attentional load in phase 3 where participants performed two tasks, as opposed to one task in phase 2.

A paired-samples *t*-test on the face task accuracy showed that participants were more accurate in reporting seeing a face in a critical trial when the face was a neutral face (*M* = 0.59, *SD* = 0.18), compared to when it was a fearful face (*M* = 0.56, *SD* = 0.16), *t*(23) = 2.84, *p* = .009, Cohen’s *d* = 0.58. The lower accuracy for the fearful face does not seem to support an attentional bias towards the fearful compared to neutral expression, as would be expected. The overall low accuracy in the face task may further suggest that the attentional load was rather high in phase 3, consequently preventing participants from readily detecting the face stimuli.

A paired-samples *t*-test showed that the d’ was higher for neutral face condition (*M* = 1.59, *SD* = 1.66) than fearful face condition (*M* = 1.53, *SD* = 1.67), *t*(23) = 2.73, *p* = .012, Cohen’s *d* = 0.56. Additionally, there seemed to be an overall response bias towards reporting not seeing a face (c = 0.92 for fearful face condition; c = 0.88 for neutral face condition).

### Mass univariate analysis

The 3(phase) x 2(laterality) x 2(face emotion) ANOVA on the difference waves between the critical and the non-critical conditions revealed a significant main effect of phase, *Fs* > 3.20, *ps* < .038. We followed up on this effect by conducting pairwise cluster-based permutation *t*-tests, using an adjusted alpha level of .017 (0.05/3).

There was no significant difference between phase 1 and 2, *ps* > .062, or between phase 2 and 3, *ps* > .064. However, when comparing phase 1 and 3, significant differences emerged, whereby the ERP signals were more positive in phase 3 than phase 1 (i.e., a LP) between 332-797ms over parietal electrodes C1/2, P1/2, P3/4 and PO3/4 (temporal peak: 633ms; spatial peak: P3/4), *ts* > 2.07, *ps* < .005, see Figure 3a & 3b.

**Figure 3.**
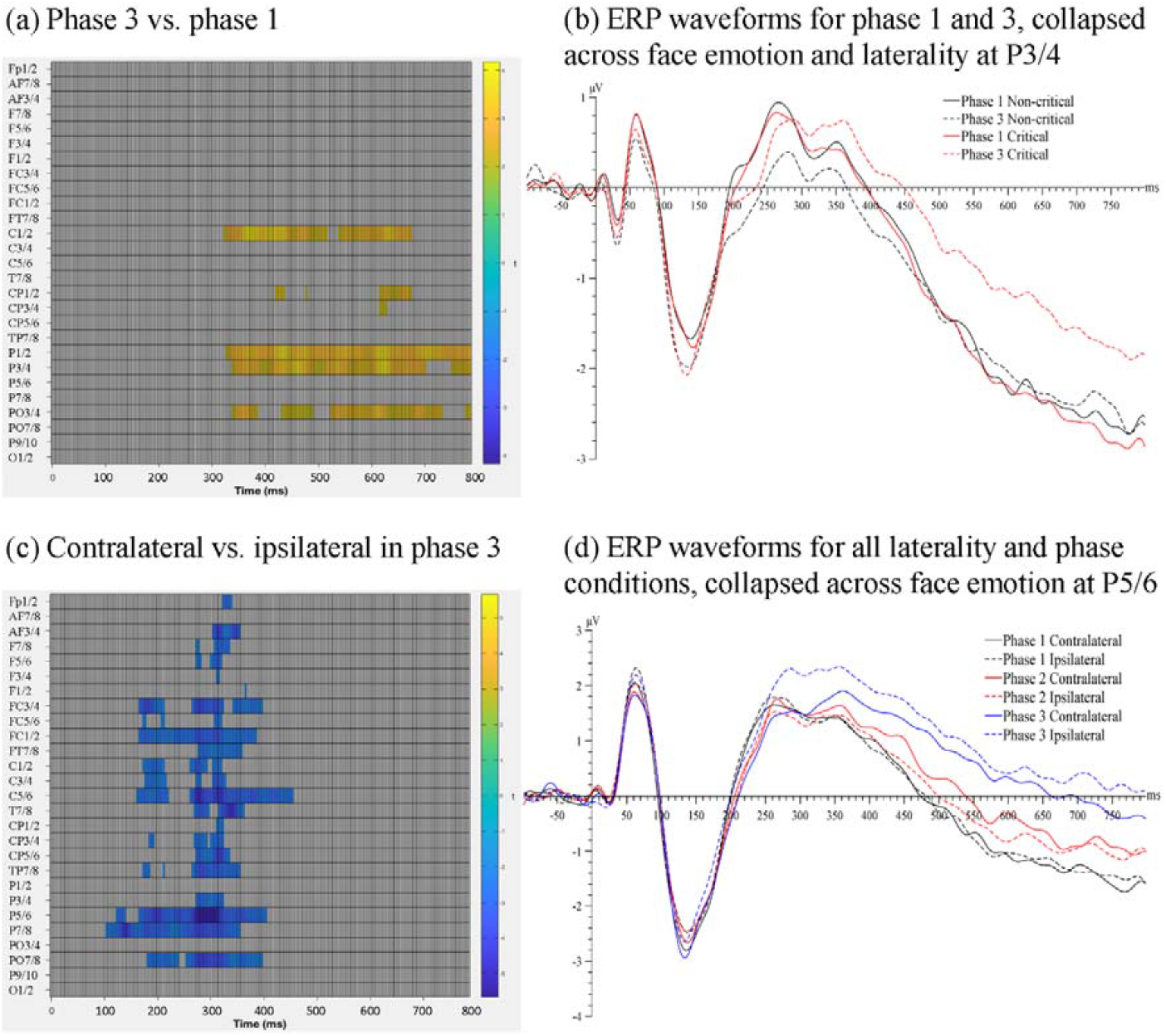
(a) Raster plot for the pairwise comparison between phase 1 and phase 3. (b) ERP waveforms for phase 1 and phase 3, collapsed across face emotion and laterality conditions, over electrodes P3/4, the pair of electrodes that showed the maximal effect. (c) Raster plot for the pairwise comparison between contralateral and ipsilateral signals in phase 3. (d) ERP waveforms for different phase and laterality conditions, collapsed across face emotion conditions, over electrodes P5/6, the pair of electrodes that showed the maximal effect.

The interaction between phase and laterality was significant, *Fs* > 3.21, *ps* < .010. To follow up this interaction, we compared contralateral and ipsilateral signals in each phase, using an adjusted alpha level of .017. The effect of laterality was not significant in either phase 1 or phase 2, *ps* > .354, but was significant in phase 3, *ts* > 2.07, *ps* < .001 (see Figure 3c). Specifically, ERP signals contralateral to the faces were more negative than ipsilateral signals between 105-410ms over several posterior electrodes (temporal peak: 285ms; spatial peak: P5/6), which reflected an N2pc to the face images (Figure 3d). That is, in the current experiment, an N2pc for faces was found only in phase 3 where participants were consciously aware of the task-relevant faces. When the participants were not aware of the faces (phase 1) or when the faces were not relevant to the task (phase 2), no N2pc was elicited by the faces.

No other omnibus effect was significant, *ps* > .085.

In an additional analysis, for phase 3, we only included trials where participants reported seeing a face and conducted a 3(phase) x 2(laterality) ANOVA, collapsed across face emotion (to increase power for this analysis). Results showed that both main effects of phase and laterality as well as the interaction effect were significant, *Fs* > 3.20, *ps* < .005. Follow-up *t*-tests showed that, an N2pc to the critical face stimulus was found in phase 3, *ts* > 2.07, *ps* < .002. Additionally, a significant posterior LP was found for phase 3, compared to both phase 1 and phase 2, *ts* > 2.07, *ps* < .005. Therefore, the phase 1-phase 3 difference found in our main analysis seems to be at least partly related to participants’ awareness of the faces.

## Discussion

In this inattentional blindness EEG study, we implemented awareness and task-relevancy of face stimuli (fearful and neutral faces) across three phases of the experiment.

Our main finding is that faces captured attention, indexed by an N2pc to the faces, only when participants were aware of the faces and when the faces were task-relevant in phase 3. This is consistent with our hypothesis and aligns with the findings from our previous backward masking studies (Qiu et al., 2022a; 2023b). It seems that, regardless of the methods used to manipulate awareness, face stimuli only attract attention when they are consciously detected and relevant for the task at hand. Importantly, we did not find any difference associated with face emotion in any phase. That is, fearful faces did not enhance the ERPs, compared to neutral faces, even when the face stimuli were task-relevant. Thus, our current results show that faces do not attract attention in an automatic manner.

There have been some observations of attention shifting towards irrelevant or unattended emotional faces, but in situations where the attentional load is thought to be rather low (Eimer & Kiss, 2007; Fox et al., 2002; Qiu et al., 2023a). For example, using a dot-probe paradigm, both Fox et al. (2002) and Qiu et al. (2023a) found that participants had an attentional bias towards task-irrelevant fearful faces. In particular, in the study by Qiu et al. (2023a), when participants were asked to localise a dot presented after a pair of masked faces, an N2pc was observed for fearful face that was clearly visible to the participants, even when faces were not relevant to the task (Qiu et al., 2023a). As we discussed, the attentional load was considered low overall in the dot-probe paradigm due to a temporal separation between the face stimuli and the target dot. As a result, spatial attention could shift to the fearful face even when it was task-irrelevant. In the present study, the accuracy for the main rectangle localisation task was only at around 80%, suggesting that the attentional load in the current IB experiment was likely higher than in these previous studies. It is likely for this reason that the task-irrelevant faces did not attract spatial attention in phase 1 and 2, as indexed by the absence of an N2pc. Similar findings have been reported by other researchers (e.g., Lien et al., 2013; Pessoa et al., 2002).

Our hypotheses about the effects of phase were not supported by the current data. First, there was no significant difference between the ERPs in phase 1 and 2. As the intended difference between these two phases was whether or not participants were aware of the face stimuli, it is surprising that we did not find a significantly enhanced negativity (a VAN) for phase 2.

In previous awareness studies, especially those using the masking techniques, the VAN has been suggested to originate from localised recurrent processes at early visual cortices and hence to reflect early perceptual awareness, as opposed to a later stage of awareness which may be indexed by the LP (Förster et al., 2020). Specifically, in a typical backward masking experiment, stimuli in the subliminal or nonconscious condition are usually presented for a very short duration (e.g., 16ms) whereas they are presented for much longer in the supraliminal or conscious condition (e.g., 266ms). Although this manipulation is integral to the backward masking technique, the visual inputs are different in these two conditions. The VAN is perhaps very sensitive to these perceptual differences and has indeed been found to robustly correlate with awareness in studies using this sensory manipulation of conscious awareness (Förster et al., 2020). In the current experiment, however, all stimuli were presented for a constant of 400ms in all three phases. As there was no perceptual difference in the stimuli *per se*, it is possible that the VAN could not readily reflect the differences between the states of awareness and unawareness. In alignment with this, previous change blindness studies showed that, when the changes between trials were small or less salient, the VAN could not be found for detected compared to undetected changes (Niedeggen et al., 2001). However, when the changes were rather obvious between trials, the VAN correlated well with change detection (Koivisto & Revonsuo, 2003).

In addition, we failed to find a significantly larger LP in phase 2 compared to phase 1. This is less surprising because the LP has been found to be associated with a variety of higher-order cognitive processes that are unrelated to awareness (for a review see Polich, 2007). Perhaps, because task requirements were the same, post-perceptual processes including the evaluative appraisal of the stimuli were the same between phase 1 and 2. Consequently, no difference on the LP could be found. Alternatively, it is possible that the main task (i.e., rectangle spatial localisation task) was rather cognitively demanding and that participants did not have sufficient spare attentional resources for the task-irrelevant stimuli in either phase 1 or 2. As a result, no difference could be observed in the ERPs between these two phases.

Second, the VAN and the LP did not show any differences when the stimuli were task-irrelevant vs. task-relevant, as there was no significant difference between phase 2 and 3 in our main analysis. We argue that phase 2 and phase 3 may be associated with different levels of attention to the critical face stimuli. Specifically, after being informed about the presence of faces, participants’ attention may be captured by the now-aware face stimulus in some but not all trials during phase 2. There is likely a certain level of fluctuation in attention across trials in phase 2, especially considering that the attentional load of the main task was high, as noted previously. By contrast, in phase 3, participants had to fully attend to the face in every trial in order to correctly perform the face task. Even when participants were required to report their awareness of the face stimuli on a trial-by-trial basis, participants failed to detect the face in a lot of critical trials (face task accuracy lower than 60%). This shows that fluctuation in attention at the trial level indeed existed throughout the experiment. The variance across participants in how much attention was directed to the faces in phase 2 may be one reason why we did not see any significant difference between the ERPs in phase 2 and 3.

Nevertheless, we found a late positivity for phase 3 when compared to phase 1. It therefore appears that participants become aware of something only when they *fully* attend to them. When not fully attended, or being task-irrelevant in this case, the faces may not be able to induce significant changes in the electrophysiological activity. This claim is consistent with a prominent view on the attention-awareness relationship which posits that attention is a prerequisite to conscious awareness (for reviews see Cohen et al., 2012; Marchetti, 2012; Noah & Mangun, 2020). Note, however, that studies favouring this argument usually use attentional paradigms such as the IB and attentional blink (Cohen et al., 2012), which are not the only methods to manipulate visual awareness. As discussed in Introduction, distinctions should be made between attentional and sensory paradigms when discussing visual (un)awareness (Diano et al., 2017). In our recent review, we have found that in attentional paradigms, unaware emotional faces were associated with stronger activation of regions in a large right-lateralised neural network that is linked to attention processes (Qiu et al., 2022c). However, this network was not found to be activated for unseen emotional faces in sensory paradigms (Qiu et al., 2022c). Perhaps, the requirement of attention before awareness is only necessary when the fate of visual information relies on attentional availability. However, when the visual inputs are made subliminal at the perceptual level, attention to the stimuli is likely futile.

Given the intricate nature of the attention-awareness relationship, we argue that studying this question requires systematic evidence gathered over all viable experimental paradigms and stimuli. Our current data add important evidence to this extensive line of research by showing that task-relevancy is necessary for attentional awareness to occur for face stimuli.

As noted, one limitation of the current study is that the main task (rectangle spatial localisation task) may have been too difficult. Participants may therefore not have sufficient attentional resources for the task-irrelevant faces in phase 2. As a result, phase 1 and phase 2 were not associated with significant ERP differences. Future studies should aim to systematically vary the task difficulty and thus, attentional load, and examine whether modulations on the awareness-related components can be found already when comparing phase 2 with phase 1.

In conclusion, in an inattentional blindness paradigm, task-relevancy of or attention to human faces seems necessary for awareness to occur. Additionally, combined with our previous work (Qiu et al., 2022a; 2023b), fearful faces or faces in general do not attract spatial attention when they are task-irrelevant.

## Acknowledgement

Zeguo Qiu was supported by Graduate Scholarships provided by University of Queensland.

## CRediT authorship contribution statement

**Zeguo Qiu:** Conceptualization, Methodology, Formal analysis, Investigation, Data curation, Writing - Original Draft, Writing – Review & Editing, Visualization, Project administration.

**Xue Lei:** Investigation. **Stefanie I. Becker**: Writing - Review & Editing. **Alan J. Pegna**: Resources, Writing - Review & Editing.

## Data availability

Data will be made available on request.

## References

Campbell, J. I., & Thompson, V. A. (2012). MorePower 6.0 for ANOVA with relational confidence intervals and Bayesian analysis. Behavior research methods, 44, 1255–1265. https://doi.org/10.3758/s13428-012-0186-0

Celeghin, A., de Gelder, B., & Tamietto, M. (2015). From affective blindsight to emotional consciousness. Consciousness and cognition, 36, 414–425. https://doi.org/10.1016/j.concog.2015.05.007

Cohen, M. A., Cavanagh, P., Chun, M. M., & Nakayama, K. (2012). The attentional requirements of consciousness. Trends in cognitive sciences, 16(8), 411–417. https://doi.org/10.1016/j.tics.2012.06.013

Cohen, M. A., Ortego, K., Kyroudis, A., & Pitts, M. (2020). Distinguishing the neural correlates of perceptual awareness and postperceptual processing. Journal of Neuroscience, 40(25), 4925–4935. https://doi.org/10.1523/JNEUROSCI.0120-20.2020

Delorme, A., & Makeig, S. (2004). EEGLAB: an open source toolbox for analysis of single-trial EEG dynamics including independent component analysis. Journal of neuroscience methods, 134(1), 9–21. https://doi.org/10.1016/j.jneumeth.2003.10.009

Del Zotto, M. D., & Pegna, A. J. (2015). Processing of masked and unmasked emotional faces under different attentional conditions: an electrophysiological investigation. Frontiers in psychology, 6, 1691. https://doi.org/10.3389/fpsyg.2015.01691

Dembski, C., Koch, C., & Pitts, M. (2021). Perceptual awareness negativity: a physiological correlate of sensory consciousness. Trends in Cognitive Sciences, 25(8), 660–670. https://doi.org/10.1016/j.tics.2021.05.009

Diano, M., Celeghin, A., Bagnis, A., & Tamietto, M. (2017). Amygdala response to emotional stimuli without awareness: facts and interpretations. Frontiers in psychology, 7, 2029. https://doi.org/10.3389/fpsyg.2016.02029

Donaldson, W. (1992). Measuring recognition memory. Journal of Experimental Psychology: General, 121(3), 275. https://psycnet.apa.org/doi/10.1037/0096-3445.121.3.275

Eimer, M., & Kiss, M. (2007). Attentional capture by task-irrelevant fearful faces is revealed by the N2pc component. Biological psychology, 74(1), 108–112. https://doi.org/10.1016/j.biopsycho.2006.06.008

Elam, K. K., Carlson, J. M., DiLalla, L. F., & Reinke, K. S. (2010). Emotional faces capture spatial attention in 5-year-old children. Evolutionary Psychology, 8(4), 147470491000800415. https://doi.org/10.1177/147470491000800415

Fields, E. C., & Kuperberg, G. R. (2020). Having your cake and eating it too: Flexibility and power with mass univariate statistics for ERP data. Psychophysiology, 57(2), e13468. https://doi.org/10.1111/psyp.13468

Förster, J., Koivisto, M., & Revonsuo, A. (2020). ERP and MEG correlates of visual consciousness: The second decade. Consciousness and cognition, 80, 102917. https://doi.org/10.1016/j.concog.2020.102917

Fox, E. (2002). Processing emotional facial expressions: The role of anxiety and awareness. Cognitive, Affective, & Behavioral Neuroscience, 2(1), 52–63. https://doi.org/10.3758/CABN.2.1.52

Fox, E., Russo, R., & Dutton, K. (2002). Attentional bias for threat: Evidence for delayed disengagement from emotional faces. Cognition & emotion, 16(3), 355–379. https://doi.org/10.1080/02699930143000527

Frischen, A., Eastwood, J. D., & Smilek, D. (2008). Visual search for faces with emotional expressions. Psychological bulletin, 134(5), 662. https://doi.org/10.1037/0033-2909.134.5.662

Glickman, M., & Lamy, D. (2018). Attentional capture by irrelevant emotional distractor faces is contingent on implicit attentional settings. Cognition and Emotion, 32(2), 303–314. https://doi.org/10.1080/02699931.2017.1301883

Groppe, D. M., Urbach, T. P., & Kutas, M. (2011). Mass univariate analysis of event□related brain potentials/fields II: Simulation studies. Psychophysiology, 48(12), 1726–1737. https://doi.org/10.1111/j.1469-8986.2011.01272.x

Grose-Fifer, J., Rodrigues, A., Hoover, S., & Zottoli, T. (2013). Attentional capture by emotional faces in adolescence. Advances in cognitive psychology, 9(2), 81. https://doi.org/10.2478/v10053-008-0134-9

Harris, A. M., Dux, P. E., & Mattingley, J. B. (2020). Awareness is related to reduced post□stimulus alpha power: a no□report inattentional blindness study. European Journal of Neuroscience, 52(11), 4411–4422. https://doi.org/10.1111/ejn.13947

Hedger, N., Garner, M., & Adams, W. J. (2019). Do emotional faces capture attention, and does this depend on awareness? Evidence from the visual probe paradigm. Journal of Experimental Psychology: Human Perception and Performance, 45(6), 790. https://doi.org/10.1037/xhp0000640

Holmes, A., Bradley, B. P., Nielsen, M. K., & Mogg, K. (2009). Attentional selectivity for emotional faces: Evidence from human electrophysiology. Psychophysiology, 46(1), 62–68. https://doi.org/10.1111/j.1469-8986.2008.00750.x

Kiss, M., & Eimer, M. (2008). ERPs reveal subliminal processing of fearful faces. Psychophysiology, 45(2), 318–326. https://doi.org/10.1111/j.1469-8986.2007.00634.x

Koivisto, M., & Revonsuo, A. (2003). An ERP study of change detection, change blindness, and visual awareness. Psychophysiology, 40(3), 423–429. https://doi.org/10.1111/1469-8986.00044

Làdavas, E., & Bertini, C. (2021). Right hemisphere dominance for unconscious emotionally salient stimuli. Brain Sciences, 11(7), 823. https://doi.org/10.3390/brainsci11070823

Langner, O., Dotsch, R., Bijlstra, G., Wigboldus, D. H., Hawk, S. T., & Van Knippenberg, A. D. (2010). Presentation and validation of the Radboud Faces Database. Cognition and emotion, 24(8), 1377–1388. https://doi.org/10.1080/02699930903485076

Lien, M. C., Taylor, R., & Ruthruff, E. (2013). Capture by fear revisited: An electrophysiological investigation. Journal of Cognitive Psychology, 25(7), 873–888. https://doi.org/10.1080/20445911.2013.833933

Lopez-Calderon, J., & Luck, S. J. (2014). ERPLAB: an open-source toolbox for the analysis of event-related potentials. Frontiers in human neuroscience, 8, 213. https://doi.org/10.3389/fnhum.2014.00213

Marchetti, G. (2012). Against the view that consciousness and attention are fully dissociable. Frontiers in psychology, 3, 36. https://doi.org/10.3389/fpsyg.2012.00036

Niedeggen, M., Wichmann, P., & Stoerig, P. (2001). Change blindness and time to consciousness. European Journal of Neuroscience, 14(10), 1719–1726. https://doi.org/10.1046/j.0953-816x.2001.01785.x

Noah, S., & Mangun, G. R. (2020). Recent evidence that attention is necessary, but not sufficient, for conscious perception. Annals of the New York Academy of Sciences, 1464(1), 52–63. https://doi.org/10.1111/nyas.14030

Pegna, A. J., Darque, A., Berrut, C., & Khateb, A. (2011). Early ERP modulation for task-irrelevant subliminal faces. Frontiers in psychology, 2, 88. https://doi.org/10.3389/fpsyg.2011.00088

Pegna, A. J., Landis, T., & Khateb, A. (2008). Electrophysiological evidence for early non-conscious processing of fearful facial expressions. International Journal of Psychophysiology, 70(2), 127–136. https://doi.org/10.1016/j.ijpsycho.2008.08.007

Peirce, J., Gray, J. R., Simpson, S., MacAskill, M., Höchenberger, R., Sogo, H., … & Lindeløv, J. K. (2019). PsychoPy2: Experiments in behavior made easy. Behavior research methods, 51, 195–203. https://doi.org/10.3758/s13428-018-01193-y

Pessoa, L., McKenna, M., Gutierrez, E., & Ungerleider, L. G. (2002). Neural processing of emotional faces requires attention. Proceedings of the National Academy of Sciences, 99(17), 11458–11463. https://doi.org/10.1073/pnas.172403899

Pitts, M. A., Lutsyshyna, L. A., & Hillyard, S. A. (2018). The relationship between attention and consciousness: an expanded taxonomy and implications for ‘no-report’paradigms. Philosophical Transactions of the Royal Society B: Biological Sciences, 373(1755), 20170348. https://doi.org/10.1098/rstb.2017.0348

Polich, J. (2007). Updating P300: an integrative theory of P3a and P3b. Clinical neurophysiology, 118(10), 2128–2148. https://doi.org/10.1016/j.clinph.2007.04.019

Pourtois, G., & Vuilleumier, P. (2006). Dynamics of emotional effects on spatial attention in the human visual cortex. Progress in brain research, 156, 67–91. https://doi.org/10.1016/S0079-6123(06)56004-2

Qiu, Z., Becker, S. I., & Pegna, A. J. (2022a). Spatial attention shifting to emotional faces is contingent on awareness and task relevancy. Cortex, 151, 30–48. https://doi.org/10.1016/j.cortex.2022.02.009

Qiu, Z., Becker, S. I., & Pegna, A. J. (2022b). Spatial attention shifting to fearful faces depends on visual awareness in attentional blink: An ERP study. Neuropsychologia, 172, 108283.

Qiu, Z., Jiang, J., Becker, S. I., & Pegna, A. J. (2023a). Attentional capture by fearful faces requires consciousness and is modulated by task-relevancy: a dot-probe EEG study. Frontiers in Neuroscience, 17:1152220. https://doi.org/10.3389/fnins.2023.1152220

Qiu, Z., Lei, X., Becker, S. I., & Pegna, A. J. (2022c). Neural activities during the Processing of unattended and unseen emotional faces: a voxel-wise Meta-analysis. Brain Imaging and Behavior, 16(5), 2426–2443. https://doi.org/10.1007/s11682-022-00697-8

Qiu, Z., Zhang, J., & Pegna, A. J. (2023b). Neural processing of lateralised task-irrelevant fearful faces under different awareness conditions. Consciousness and Cognition, 107, 103449. https://doi.org/10.1016/j.concog.2022.103449

Reynolds, G. D., & Roth, K. C. (2018). The development of attentional biases for faces in infancy: A developmental systems perspective. Frontiers in psychology, 9, 222. https://doi.org/10.3389/fpsyg.2018.00222

Suzuki, M., & Noguchi, Y. (2013). Reversal of the face-inversion effect in N170 under unconscious visual processing. Neuropsychologia, 51(3), 400–409. https://doi.org/10.1016/j.neuropsychologia.2012.11.021

Tipura, E., & Pegna, A. J. (2022). Subliminal emotional faces do not capture attention under high attentional load in a randomized trial presentation. Visual Cognition, 30(4), 280–288. https://doi.org/10.1080/13506285.2022.2060397

Tsuchiya, N., Wilke, M., Frässle, S., & Lamme, V. A. (2015). No-report paradigms: extracting the true neural correlates of consciousness. Trends in cognitive sciences, 19(12), 757–770. https://doi.org/10.1016/j.tics.2015.10.002

Zhou, L. F., Wang, K., He, L., & Meng, M. (2021). Twofold advantages of face processing with or without visual awareness. Journal of Experimental Psychology: Human Perception and Performance, 47(6), 784. https://doi.org/10.1037/xhp0000915

